# KIN-29 SIK regulates stress-induced sleep through mitochondrial redox signaling

**DOI:** 10.64898/2026.07.27.740804

**Authors:** Pearson McIntire, Laura N. Farrell, Suraiya Haroon, David M. Raizen, Alexander M. van der Linden

## Abstract

The *C. elegans* SIK3 homolog KIN-29 regulates the interaction between sleep and metabolism, but mechanisms underlying this regulation are not understood. Here, we show that KIN-29 regulates sleep that is induced by cellular stress (stress-induced sleep or SIS) through mitochondrial reactive oxygen species (ROS) signaling. Following sleep-promoting ultraviolet-C (UVC) irradiation, mitochondrial ROS rises in concert with sleep in wild-type but not in sleepless *kin-29* mutants. *kin-29* mutants have reduced mitochondrial ROS, reduced oxygen consumption rates, and are resistant to oxidative stress. Transcriptomic and proteomic profiling of *kin-29* mutants reveal enrichment for genes involved in ROS mitigation such as the mitochondrial superoxide dismutase SOD-3. Consistent with the notion that ROS promotes sleep, genetic disruption of mitochondrial SODs enhances UVC-induced SIS. Increasing mitochondrial ROS using the optogenetic tool SuperNova partially restores SIS in *kin-29* mutants but not in mutants with defective function of the sleep-inducing ALA neuron; this suggests that mitochondrial ROS act downstream of *kin-29* but upstream of ALA. The identification of mitochondrial ROS as a nematode sleep regulator supports a phylogenetically conserved mechanism by which metabolic stress and SIKs promote sleep.

## INTRODUCTION

Salt-inducible kinases are evolutionarily conserved regulators of both sleep and metabolism (*1*). A mouse forward genetic screen identified a gain-of-function *Sik3* allele that increases sleep duration and intensity (*2–5*). Reduced SIK3 activity is associated with reduced sleep both in mice (*6*) and in humans (*7*). These studies establish SIK3 as a conserved regulator of sleep.

KIN-29, the sole *Caenorhabditis elegans* homolog of the three mammalian SIK genes provides a genetically tractable model for understanding how this kinase couples metabolism to sleep. Loss of *kin-29* in glutamatergic sensory neurons reduces sleep and increases body fat; the effect of *kin-29* on sleep occurs upstream of the regulation of the sleep-promoting ALA and RIS neurons (*8*). These findings suggest that KIN-29 links metabolic information to sleep-regulatory circuits. However, the signal that conveys this metabolic state to sleep-promoting neurons remains unclear.

One candidate is mitochondrial redox state signals. In *C. elegans*, inhibition of mitochondrial β-oxidation, a major source of reactive oxygen species (ROS), increases fat accumulation (*9*) and reduces stress-induced sleep (*8*). This raises the possibility that ROS are not merely metabolic byproducts, but sleep-promoting signals downstream of KIN-29. Endogenous ROS, and in particular hydrogen peroxide (H₂O₂), regulate cellular homeostasis, while excessive ROS increases shift an organism’s redox tone to oxidative distress. Thus, KIN-29 promotion of ROS may bridge metabolism, cellular stress, and sleep.

Interactions between ROS and sleep occurs in both insects and mammals (*10*). In *Drosophila*, short-sleeping mutants are sensitive to oxidative stress, increasing sleep improves survival after oxidative challenge, and neuronal antioxidant overexpression reduces sleep, supporting a bidirectional relationship between sleep and oxidative stress (*11*). Sleep loss elevates mitochondrial ROS in dorsal fan-shaped body (dFB) neurons of *Drosophila*, where a potassium channel subunit is proposed to sense the redox-state of the animal and thus increase neuronal excitability and promotes sleep (*12, 13*). In mice, cytosolic H_2_O_2_ accumulates in sleep-promoting mid-brain regions during extended wakefulness, track sleep pressure and promoted sleep initiation within a physiological ROS range (*10, 14*). Together, these studies suggest that redox signals can encode sleep need and promote sleep.

ROS also links sleep to systemic physiology. Severe sleep deprivation causes ROS accumulation in the gut of flies and mice, and preventing ROS accumulation through antioxidants or gut-targeted antioxidant enzymes like Superoxide Dismutase (SOD) allows sleep-deprived flies to survive with little or no sleep (*15, 16*). Moreover, sleep deprivation in zebrafish induces gut ROS accumulation (*17*). Thus, ROS may act at multiple levels: as circuit level signals that promote sleep, and as systemic mediators of sleep loss.

Key questions remain: does the role of ROS signaling in sleep regulation extend across phylogeny beyond vertebrates and arthropods? Is ROS signaling altered by key sleep genes such as salt inducible kinases?

*C. elegans* provides a powerful model system to answering these questions. Stress-induced sleep (SIS) is triggered by environmental insults, which include heat shock (*18*), infection (*19, 20*), mechanical tissue damage (*21, 22*), toxin exposure (*18*), and ultraviolet radiation exposure (*23*). This state is characterized by behavioral quiescence, reduced feeding, and elevated arousal thresholds, and shares key features with sickness-associated sleep in mammals and *Drosophila* (*24*). Because *C. elegans* has a conserved SIK homolog, KIN-29, and undergoes robust SIS in response to stressors, it offers an opportunity to test whether ROS-dependent sleep regulation extends to nematodes and whether KIN-29/SIK links metabolic stress to ROS-driven sleep.

Here, we investigated the role of ROS in KIN-29-mediated regulation of SIS. We find that KIN-29 promotes SIS through mitochondrial redox signaling. *kin-29* mutants show reduced levels of mitochondrial ROS, lower basal respiration, reduced respiration response to mitochondrial stress, and defective mitochondrial ROS induction in response to UVC-irradiation. Transcriptomic and proteomic analyses of *kin-29* mutants reveal changes in genes involved in ROS stress responses such as *sod-3*. Animals harboring loss-of-function mutations in the mitochondrial sods or in all five known *C. elegans sods* increases UVC-induced SIS. Optogenetic induction of mitochondrial ROS by SuperNova partially restores sleep to *kin-29* mutants, showing that ROS promotes sleep downstream of *kin-29* function. In contrast, promotion of sleep by mitochondrial ROS is blocked in a mutant with dysfunction of a SIS-promoting ALA neuron, suggesting that mitochondrial ROS acts within or upstream of ALA. Together, these findings identify mitochondrial redox signaling as a downstream effector of KIN-29 and reveal a conserved mechanism through which metabolic cellular stress is translated into sleep.

## RESULTS

### Reduced ROS and enhanced oxidative stress resistance in *kin-29* mutants

We examined total ROS levels between *kin-29* mutants and wild-type worms using the 2’,7’-dichlorodihydrofluorescein diacetate (H_2_DCF-DA) probe, a dye that diffuses into cells and is oxidized by ROS to generate the fluorescent product DCF (*25*). After normalizing to total protein levels, DCF-based fluorescence levels were significantly lower in lysates from *kin-29* mutants than from wild-type controls (**Fig. 1A**). If *kin-29* mutants have reduced endogenous ROS levels, then we would expect *kin-29* mutants to be less sensitive to exogenous ROS. To test this prediction, we assessed the sensitivity of adult *kin-29* mutants to exogenous oxidative stress using the ROS-inducing agent paraquat (PQ). As a positive control for partial PQ resistance, we found increased survival in *flcn-1* mutants, which have previously been reported to be PQ-resistant (*26*) (**Fig. 1B**). Like *flcn-1* mutants, in comparison to wild-type controls, *kin-29* mutants were resistant to the lethal effects of 100 mM PQ (**Fig. 1B**). These findings show that loss of KIN-29 results in reduced ROS and reduced sensitivity to oxidative stress.

**Fig. 1.**
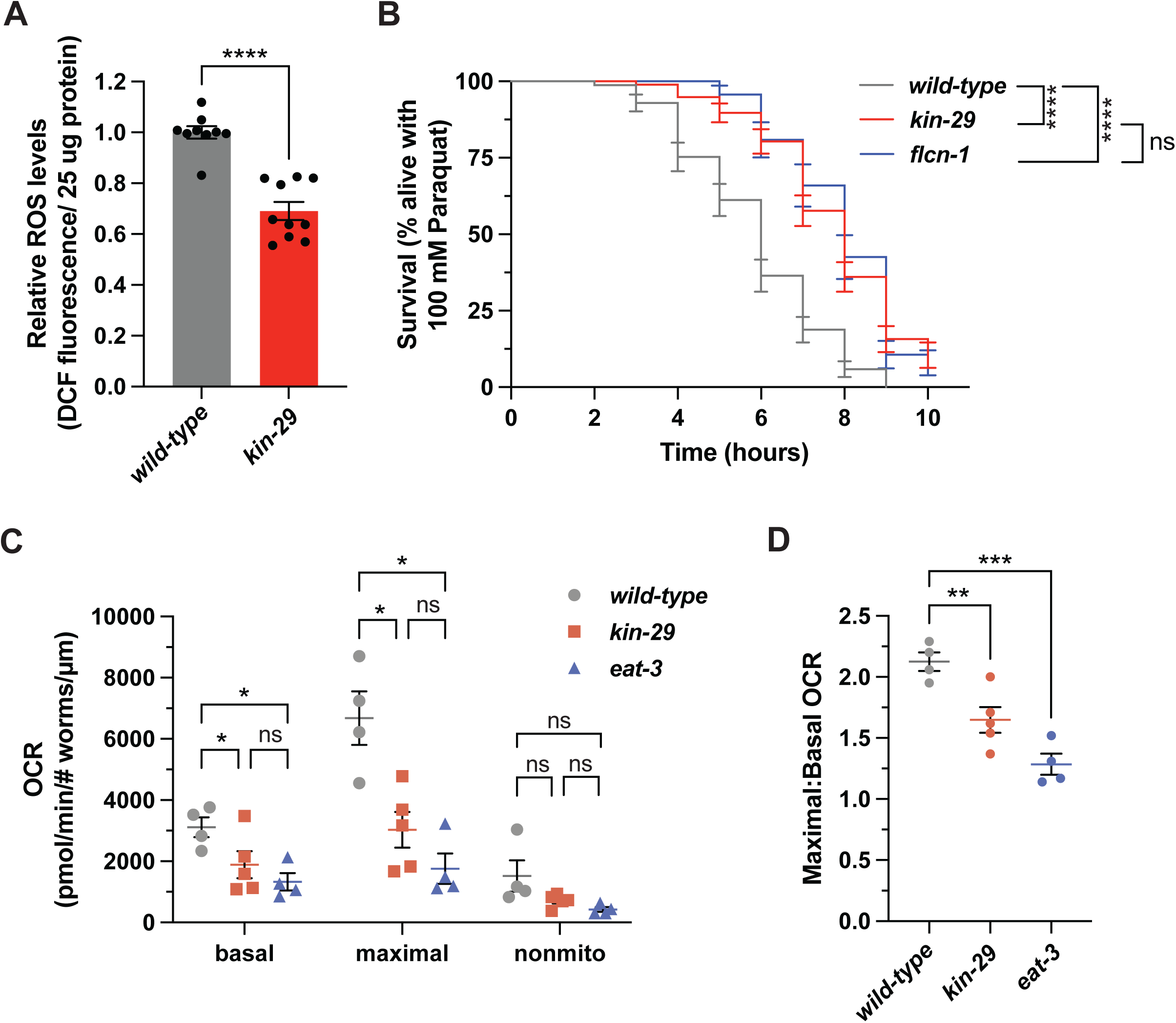
*kin-29* mutants have reduced basal ROS and paraquat resistance. **(A)** Total intracellular ROS levels per 25 ug protein in awake wild-type and *kin-29(oy38)* mutants. ROS was measured by staining whole animal lysates with 25 μM H_2_DCFDA. Data are normalized to wild-type (*wt*) and represented as the mean ± SEM of 3 biological replicates, each was which is the average of 3-4 technical replicates. \*\*\*\**P*<0.0001 by a Mann-Whitney test. **(B)** Acute paraquat exposure survival assay showing the percentage of animals alive at each time-point. n=3 biological trials for each genotype with >25 animals per trial. *flcn-1(ok975)* mutants were used as a control for resistance to paraquat (*26*). Lifespan curves were compared using the Log rank Mantel-Cox test followed by a Holm-Sidak’s multiple comparisons test; \*\*\*\**P*<0.0001; ns, not significant. **(C)** Oxygen Consumption Rate (OCR) measurements of wild-type (*wt*), *kin-29(oy38)*, and *eat-3(ad426)* mutant L4s during basal conditions, and treated with the uncoupler FCCP (25 uM) to determine maximal OCR, and the complex IV inhibitor NaN_3_ (50 mM) to determine non-mitochondrial OCR using a Seahorse XF Analyzer. OCR readings were normalized by size and number of worms per well for each genotype. Data are represented as the mean ± SEM. *wild-type* (n=4), *kin-29* (n=5), and *eat-3* (n=4) biological replicates. Significance was assessed using a Mixed-effects analysis followed by Tukey’s multiple comparisons test; \**P*<0.05. Basal = basal mitochondrial respiration, Maximal = maximal respiration, Non-mito = non-mitochondrial respiration. **(D)** Maximal:basal OCR ratio of each experimental replicate. Data are represented as the mean ± SEM. Significance was assessed using a one-way ANOVA followed by Dunnett’s multiple comparisons test; \*\*\**P*<0.001, \*\**P*<0.01.

Reduced ROS could be explained by reduced oxidative phosphorylation via the electron transport chain (ETC). To test ETC function, we measured the oxygen consumption rate (OCR) of *kin-29* mutants using a Seahorse XF analyzer, where *eat-3(ad426)* animals known to have ETC defects (*27*), are used as a control. Since *kin-29* and *eat-3* mutants are smaller than wild-type animals (*28, 29*), we normalized OCR measurements to body size as well as to animal number. Our analysis showed that the basal respiration rate of *kin-29* mutants is reduced in comparison to the basal respiration rate of wild-type worms and was not significantly different from that of *eat-3* mutants **(Fig. 1C-D, Fig. S1)**.

To measure the maximal mitochondrial capacity of the worms, we treated the animals with carbonyl cyanide-p-(trifluoromethoxy)phenylhydrazone (FCCP), which mimics a physiological maximal energy demand, stimulating the respiratory chain to operate at maximum capacity (*30*). Uncoupling the mitochondria’s proton gradient using FCCP showed that the *kin-29* mutants had a reduced response to maximal stress (**Fig. 1C-D, Fig. S1**). The FCCP-stimulated OCR of *eat-3* mutants was similarly reduced and did not differ significantly from that of *kin-29* mutants (**Fig. 1C-D, Fig. S1**). Addition of sodium azide disrupts mitochondrial oxygen consumption, which showed that there was no difference in non-mitochondrial oxygen consumption among wild-type, *kin-29* and *eat-3* mutant animals (**Fig. 1C-D, Fig. S1**). Together with their low ROS levels, these findings indicate that *kin-29* mutants exist in a reduced oxidative state and have a diminished response to higher energy demands during stress.

### KIN-29 is necessary for mitochondrial ROS increases during SIS

Our observation that *kin-29* mutants exhibit both reduced total ROS levels and reduced SIS leads to the hypothesis that ROS promotes SIS. This hypothesis predicts that ROS levels should correlate with sleep behavior in longitudinal observations. To test this, we measured ROS at three timepoints following UVC irradiation. We measured total ROS levels in wild-type animals prior to maximal sleep (0.5 and 1.0 hours post-UVC) as well as during maximal sleep (2 hours post-UVC). Following UVC irradiation (254 nM, 1,500 J/m^2^), and measured ROS using the H_2_DCF-DA fluorescence assay normalized to total protein. ROS levels were significantly elevated at 2 hours but not at 30 or 60 minutes after UVC exposure (**Fig. 2A**), supporting a close association between ROS and sleep. The notion that the ROS is causal to sleep and not simply associated with sleep is discussed further below in the section describing Supernova optogenetic experiments. In contrast to wild-type animals, *kin-29* mutant animals showed a blunted ROS response to UVC irradiation (**Fig. 2A**). Although total ROS levels did increase in *kin-29* mutants by the 2-hour time point post UVC exposure, the magnitude of this increase was lower than that observed in wild-type worms during the period when sleep is normally initiated.

**Fig. 2.**
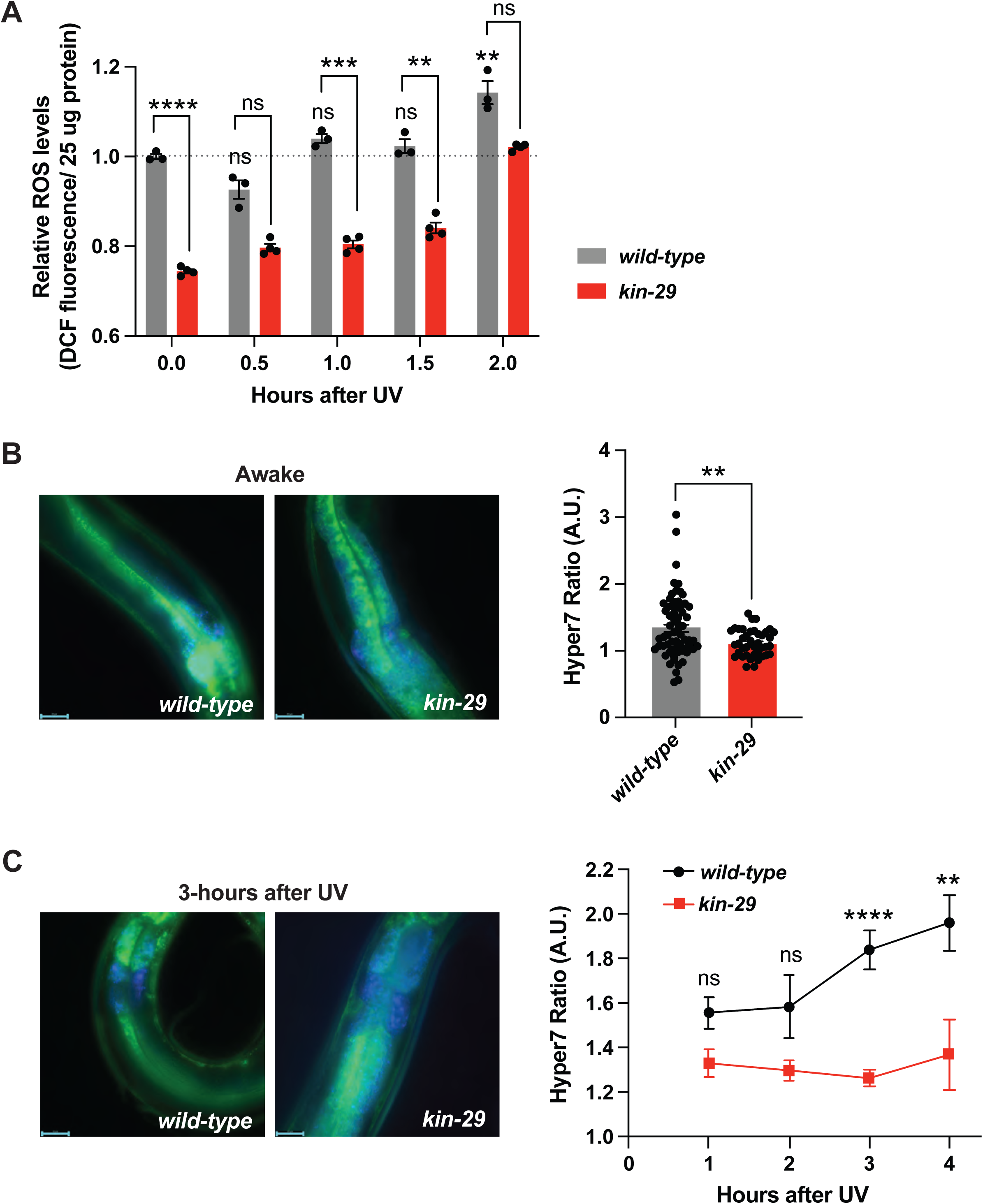
*kin-29* is necessary for mitochondrial ROS elevation during SIS. **(A)** Total ROS levels per 25 ug protein in wild-type and *kin-29(oy38)* mutants during SIS after exposure to UVC. ROS was measured by staining whole animal lysates with 25 μM H_2_DCFDA in 30 min intervals during the first 2-hours. Data were normalized to wild-type (0 hour) and significance between time-points was assessed using a 2-way ANOVA with Tukey multiple comparisons test. Data are represented as the mean ± SEM of 3-4 technical replicates for each time-point. Statistical comparisons were also performed with a 2-way ANOVA using time and genotype (wild-type and *kin-29*) as factors, followed by Bonferroni’s multiple comparisons test. \*\*\*\**P*<0.0001, \*\*\**P*<0.001 and \*\**P*<0.01; ns, not significant. **(B)** Right panel: Representative images of awake wild-type and *kin-29* mutants carrying the HyPer7 transgene. Scale bar is 25 μm. Left panel: Comparison of mitochondrial ROS during wakefulness between wild-type and *kin-29(oy38)* mutants. Quantification of fluorescence of an intestinal mitochondrial-matrix targeted HyPer7 (*Peft-3::2xCox8a::HyPer7*) (*31*). HyPer7 ratio (500/400 nm) images were obtained to measure H_2_O_2_ levels (see material and methods). Data are represented as the mean ± SEM. \*\**P*<0.01 by a two-tailed Mann-Whitney test. **(C)** Left panel: Representative images of wild-type and *kin-29(oy38)* mutants animals carrying the HyPer7 transgene 3-hours after UVC exposure. Scale bar is 25 μm. Right panel: Quantification of fluorescence of HyPer7 during SIS in wild-type (*wt*) and *kin-29* mutants following UVC exposure. HyPer7 ratio (500/400 nm) images were obtained to measure H_2_O_2_ levels (see material and methods). Data are represented as the mean ± SEM with n ≥ 7 animals for each time-point and genotype. Statistical comparisons were performed with mixed effects analysis using time and genotype as factors, followed by post-hoc comparisons at each timepoint to obtain nominal *P* values. Nominal *P* values were then subjected to Bonferroni corrections to account for multiple comparisons; \*\*\*\**P*<0.0001, \*\**P*<0.01; ns, not significant.

To assess mitochondrial ROS *in vivo* during wakefulness and SIS, we employed the mitochondria matrix-targeted HyPer7 biosensor (*31*). HyPer7 is a pH-stable reporter of H₂O₂ whose excitation maximum shifts from 400 nm to 500 nm upon oxidation (*32*). Using a compound microscope, we measured HyPer7 fluorescence ratios (excitation at 500 nm divided by excitation at 400 nm) from individual animals during wake as well as during UVC-induced SIS. HyPer7 transgenic animals exhibited a normal SIS response after UVC (**Fig. S2**), indicating that expression of the reporter does not disrupt sleep induction by stress. *kin-29* mutants exhibited lower mitochondrial ROS as measured by HyPer7 both during wakefulness (**Fig. 2B)** and throughout the 3-4 hour post-UVC window in which SIS is most robust (**Fig. 2C**). In contrast to *kin-29* mutants, wild-type animals displayed clear increases in mitochondrial ROS during hours 3-4 of SIS, whereas *kin-29* mutants failed to generate a comparable mitochondrial ROS response (**Fig. 2C**). Together, these findings suggest that KIN-29 is required for the normal rise in mitochondrial ROS during SIS.

### Transcriptomic and proteomic profiling identifies altered SOD and stress response pathways in *kin-29* mutants

Given that *kin-29* mutants exhibit reduced mitochondrial ROS during SIS and increased resistance to oxidative stress, we sought to identify the molecular pathways mis-regulated in the absence of KIN-29. We employed transcriptomic and proteomic analysis, comparing *kin-29* mutants to wild-type worms.

To identify KIN-29-dependent changes at the developmental stage in which SIS is assessed, we performed the profiling experiment on day 1 adults. We compared mRNA and protein abundance in *kin-29(oy38)* mutants to those of wild-type animals (**Fig. 1A**). RNA-seq analysis (5 biological replicates per genotype) identified 1,847 differentially expressed transcripts (|log₂ fold change| >1.5, FDR-adjusted *P*<0.05), with 1,796 upregulated and 51 downregulated genes in *kin-29* mutants (**Fig. 3B, Table S3**).

**Fig. 3.**
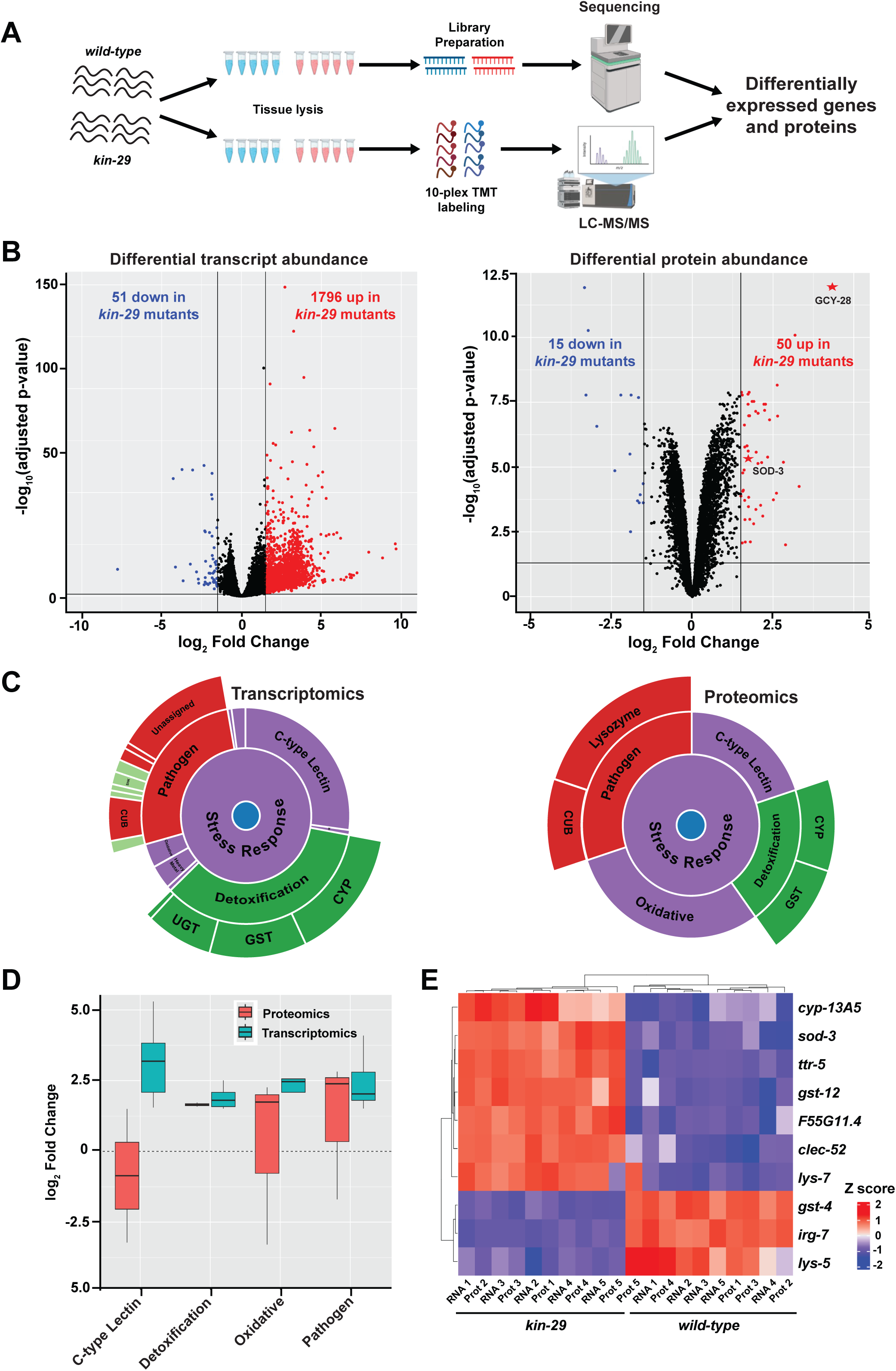
Identification of transcriptional and protein targets of KIN-29. **(A)** Schematic of collection methods for transcriptomics and proteomics. **(B)** Volcano plot of log_2_-transformed fold change against the negative log_10_ adjusted *P*-value generated using the R-package ggplot2 (*62*). Shown are the volcano plots of differentially expressed transcripts (left plot) and differentially expressed proteins (right plot). The vertical black lines represent the log_2_ Fold Change >1.5 and the horizontal black line denotes the FDR adjusted *P*-value cutoff of 0.05. Transcripts that significantly (Benjamini-Hochberg adjusted *P* value<0.05) increase or decrease in *kin-29* mutants are shown in red or blue, respectively. **(C)** WormCat 2.0 sunburst (*63*) showing key stress response subcategories for differentially expressed transcripts (left panel) and differentially expressed proteins (right panel). **(D)** Box plot showing log_2_ fold changes of genes from key stress response subcategories in the differentially abundant proteins (red bars) and differential expressed transcripts (blue bars). A positive log_2_ fold change value represents a gene that is increased in *kin-29(oy38)* mutants and a negative value is decreased in *kin-29(oy38)* relative to wild-type. **(E)** Heat map depicting the expression of all shared genes in the stress response category by sample for both omics platforms. Individual proteomics samples are denoted “prot N” and individual transcriptomics samples are denoted “RNA N”, where “N” represents a number between 1 and 5.

LC-MS/MS-based quantitative proteomic analysis (5 biological replicates per genotype) using the same statistical criteria as those used for transcriptomics, identified 65 differentially expressed proteins (out of 5,595 total proteins detected), including 50 upregulated and 15 downregulated in *kin-29* mutants relative to wild-type controls (**Fig. 3B, Table S4**). Principal component analysis (PC) showed a high degree of expression similarity among protein as well as transcripts of the same genotype, and showed clear separation between genotypes (**Fig. S3A-B**). The observation that the number of up-regulated genes and proteins was much higher than the number of down-regulated genes and proteins, indicates that KIN-29 functions primarily as a negative regulator of gene and protein expression.

To determine whether particular biological processes were enriched among the differentially expressed transcripts and proteins, we performed functional category and GO term enrichment analysis using the WormCat 2.0 (*33*) tool. A key category that was enriched in both the differentially expressed proteins and transcripts was stress response pathways for which the subcategories: oxidative, pathogen, detoxification, and C-type lectins were shared in both omics platforms (**Fig. 3C, Table S5).** Gene expression for genes associated with detoxification, oxidative, and pathogen-associated subcategories showed changes with the same directionality across both transcriptomic and proteomic data, while C-type lectins changed in the opposite direction (**Fig. 3D**). Out of the stress response genes that were identified by both platforms, the majority had the same directionality of expression including increases in expression of *cyp-13A5* (cytochrome P450), *lys-7* (lysozyme-like antimicrobial effectors), and *sod-3* in *kin-29* mutants (**Fig. 3E**)

At the mRNA level, many of the *gst* (glutathione S-transferase) gene *C. elegans* family, which contains more than 50 genes (*34*), *kin-29* mutants were upregulated in *gst* genes *kin-29* mutants: 13 *gst* genes *kin-29* mutants were upregulated and 2 were down-regulated in *kin-29* mutants (**Fig. 3E**, **Table S3**). At the protein level, changes were more modest: there was one GST protein (GST-12) upregulated and one (GST-4) down-regulated (**Table S4**).

Notably, *sod-3*, which encodes a Fe/Mn-dependent super oxide dismutase (*35*) was elevated at both the transcript and protein level in *kin-29* mutants (**Fig. 3B** and **3E**). A *Psod-3::GFP* reporter showed elevated fluorescence in *kin-29* mutants, suggesting that the *sod-3* up-regulation of mRNA in *kin-29* mutants can be attributed to the *sod-3* promoter, rather than due to reduced degradation of the transcript (**Fig. S3C-D**). The increased *sod-3* expression may partially explain the observations of reduced ROS in *kin-29* mutants (**Fig. 1**).

In summary, the multi-omics analysis reveals that *kin-29* mutation induces widespread regulation of stress-related pathways, with particularly strong enrichment for detoxification enzymes, oxidative-stress regulators, and SOD-family components. These changes are consistent with the increased oxidative stress resistance, reduced mitochondrial ROS, and reduced maximal oxygen consumption rates observed in *kin-29* mutants.

### Loss of mitochondrial SODs amplifies the somnogenic effects of UV stress

Since *sod-3* transcript and protein expression were elevated in *kin-29* mutants, we investigated whether loss of individual *sod* genes modulates SIS. ROS have been implicated as regulators of sleep in *Drosophila* and mice with SOD activity playing a central role in maintaining cellular redox balance by detoxifying superoxide radicals, a form of ROS (*11, 15*). *C. elegans* encodes five *sod* genes with partially overlapping functions (*36*). SOD-1 and SOD-5 are primarily cytoplasmic Cu/Zn-SODs (*37, 38*), SOD-2 and SOD-3 are mitochondrial Fe/Mn-dependent SODs (*35, 39*), and SOD-4 is proposed to function extracellularly or at the cell surface (*40*).

Among the five sods, *sod-3* and *sod-5* are transcriptionally induced under stress conditions (*41, 42*), suggesting roles in adaptive oxidative stress responses. Our mRNA profiling experiment revealed a 5.9-fold increase in *sod-3*, and a 4.2 elevation of *sod-5* in *kin-29* mutants (**Table S3**). Our proteomics experiment revealed a 3.5-fold elevation of SOD-3 but whether or not SOD-5 protein is differentially expressed remains an unknown because it was not among the 5,995 detected proteins (**Table S4**).

Mutants lacking mitochondrial *sod-3* displayed a modest but significant increase in movement quiescence during SIS (**Fig. 4A**), whereas two independent alleles of the mitochondrial *sod-2* showed no detectable SIS change in the first 4 hours after UVC exposure when maximal sleep occurs (**Fig. 4B-C**). However, combining *sod-2* and *sod-3* mutations further enhanced the increased SIS phenotype during the first 4 hours relative to *sod-3* or *sod-2* single mutants (**Fig. 4D**), indicating partial redundancy between mitochondrial SODs.

**Fig. 4.**
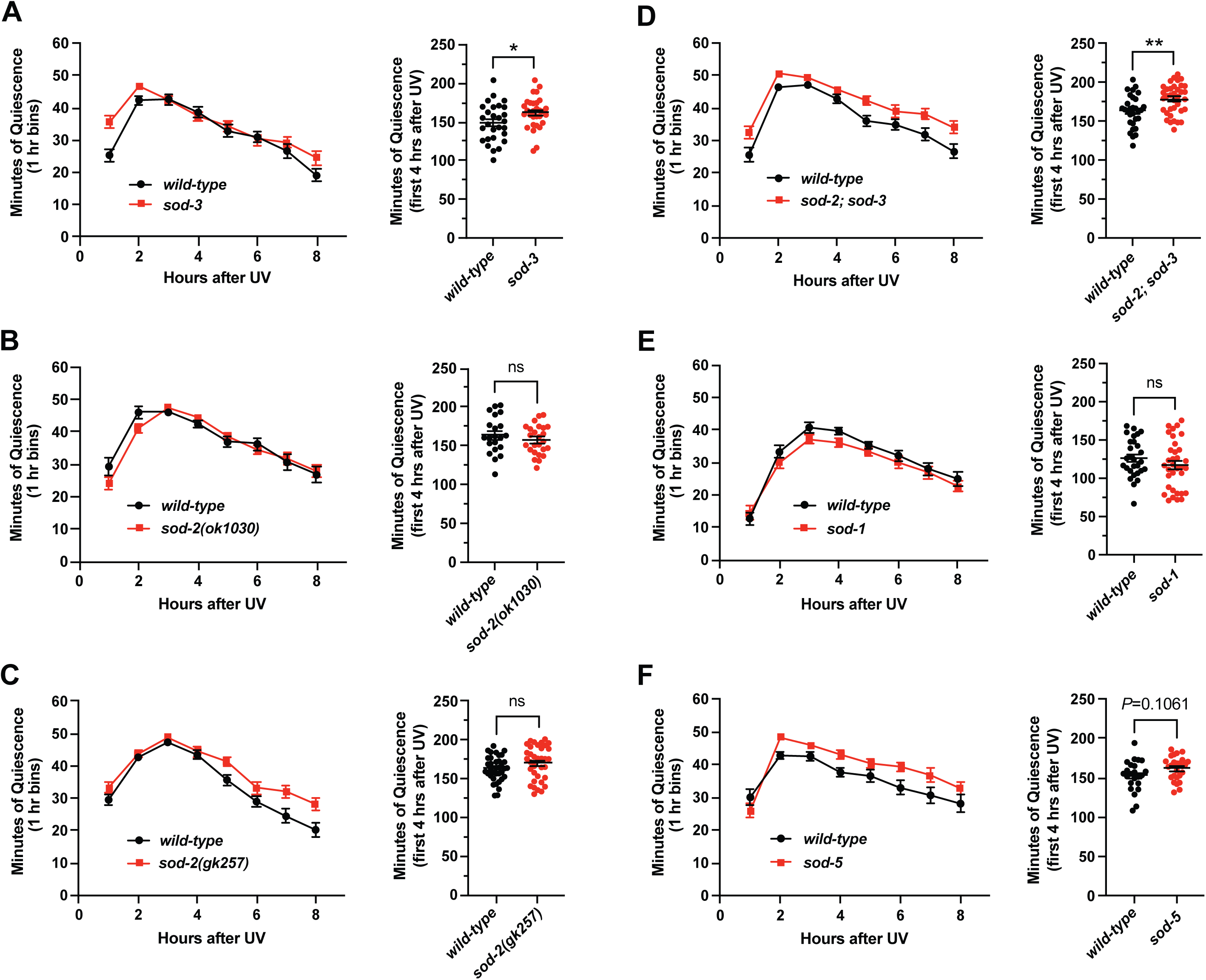
Loss of mitochondrial *sod* genes increases SIS. Time-course of body movement quiescence measured hourly over an 8-hour period (left graph) and minutes of quiescence during the first 4-hours (right graph) after UV-C exposure for **(A)** *sod-3(tm760),* **(B)** *sod-2(ok1030),* **(C)** *sod-2(gk257)*, **(D)** *sod-2(gk257);sod-3(tm760)* double mutants, **(E)** *sod-1(tm776)*, and **(F)** *sod-5(tm1146)* mutants compared to wild-type (*wt*) controls. n≥21 animals for each genotype after UV-C exposure (1,500 J/m^2^). Data are represented as the mean ± SEM of 3 trials for each genotype. Statistical comparisons were performed with a Mann-Whitney test **(C)** or by a Welch’s *t*-test **(A-B;D-F)**; \*\**P*<0.01, \**P*<0.05; ns, not significant.

In contrast to the mitochondrial *sod-3*, mutations in the cytosolic *sod-1* or *sod-5* (**Fig. 4E-F)**, or in the extracellular/membrane-associated *sod-4* (**Fig. S4**), did not significantly alter SIS in the four hours after UVC exposure. The strongest SIS phenotype was observed in the quintuple *sod* mutant lacking all five *sod* genes (*36*) at a standard UVC irradiation dose (**Fig. 5A**); comparing the phenotype of the quintuple mutant to the single *sod-3* mutant suggests that the other 4 sods may partially compensate for the loss of *sod-3* alone. We tried but were unable to create a CRISPR/Cas9 mutation in *kin-29* within the quintuple *sod* mutant, presumably because such animals were not viable. Therefore, whether the *kin-29* SIS phenotype can be attributed fully to elevated SOD activity remains unknown.

**Fig. 5.**
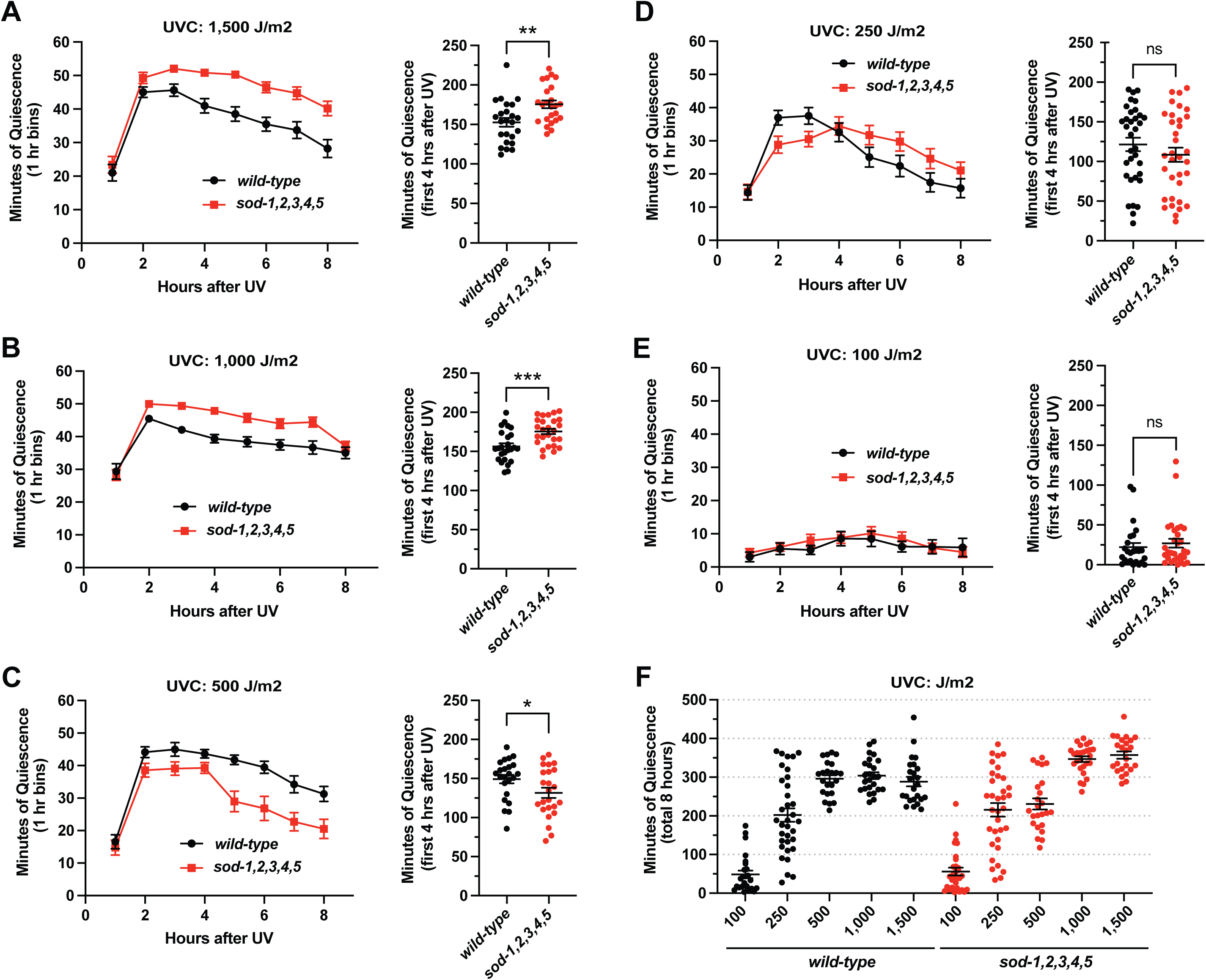
Increased SIS observed in the *sod* quintuple mutant is UV dose dependent. Time-course of body movement quiescence measured hourly over an 8-hour period (left graph) and minutes of quiescence during the first 4-hours (right graph) for the *sod*- *1,2,3,4,5* quintuple mutant after the following dosages of UV-C irradiation: **(A)** 1,500 J/m^2^, **(B)** 100 J/m^2^, **(C)** 250 J/m^2^ **(D)** 500 J/m^2^ and **(E)** 1,000 J/m^2^ dose. n≥23 animals for each genotype after UV-C irradiation. Data are represented as the mean ± SEM of 3 trials for each UV-C dosage. Statistical comparisons were performed with a Mann-Whitney test **(A-C)** or by a Welch’s *t*-test **(D-E)**; \*\*\**P*<0.001, \*\**P*<0.01, and \**P*<0.05; ns, not significant. **(F)** Summary of the total 8-hours of movement quiescence for each UV-C dosage visualized in a scatterplot.

Together, these results indicate that defects in SOD-mediated ROS clearance promotes SIS following UV stress. The enhanced SIS observed upon loss of mitochondrial SODs, particularly in the *sod-2;sod-3* double mutant, suggests that mitochondrial ROS may contribute to the somnogenic effects of UV stress and may act downstream of KIN-29 to regulate SIS.

### UV dose-dependent modulation of SIS in *sod* quintuple mutants

A possible interpretation of the *sod* quintuple mutant exhibiting increased SIS is that these mutants are generally locomotion-defective due to a combination of genetic manipulations or from chronic oxidative stress and therefore move less in response to UVC irradiation. If this interpretation is correct, then we should observe increased movement quiescence not only at the high UVC dose (1500 J/m2), but also at lower UVC dosages, when there is little to SIS. To test for this possibility, we performed SIS assays in *sod* quintuple mutants while systematically reducing the dose of UVC irradiation (254 nM) dosage from the standard 1,500 to progressively lower levels (1,000, 500, 250 and 100 J/m^2^). At high UVC doses (1,500 and 1,000 J/m^2^), *sod* quintuple mutants exhibited significantly increased movement quiescence compared to wild-type during the 4-hours following UVC irradiation (**Fig. 5A-B**). At intermediate UVC dose of 500 J/m^2^, this difference was markedly reduced with a small decrease in quiescence in the *sod* quintuple mutant compared to wild-type (*P*=0.044) (**Fig. 5C**). At lower UVC doses (250 and 100 J/m^2^), *sod* quintuple mutants displayed SIS responses that were indistinguishable from wild-type, both in the hourly quiescence time-course and during the first 4-hours after UVC exposure (**Fig. 5D-E**). Analysis of total quiescence across the post-irradiation 8-hour period further showed that quintuple *sod* mutants were disproportionally affected at higher UVC doses indicating a UV-dose dependent increase in SIS. Together, these data show that reducing the magnitude of the oxidative insult is sufficient to restore the elevated SIS in quintuple *sod* mutants to wild-type levels. This result supports the interpretation that increased SIS at high UVC doses observed in quintuple *sod* mutants is not due to non-specific generalized locomotory defects, but rather it reflects an impaired ability to mitigate ROS-induced stress.

### Inducing mitochondrial ROS is sufficient to promote SIS and partially restore SIS in *kin-29* mutant animals

The observation of a correlation between ROS levels and SIS does not prove the ROS is causing the sleep. An alternative is that ROS may rise as a consequence of sleep, or that ROS increases in parallel to sleep increase. To distinguish between these possibilities, we tested whether inducing ROS experimentally can promote sleep in the absence of UVC light. We used transgenic animals expressing the optically active ROS-generating enzyme Supernova targeted to the Mitochondrial matrix Complex I (MCI) (*31*), which is a major site of mitochondrial ROS production (**Fig. 6A**). We generated ROS by exposing animals carrying MCI-Supernova to 580 nm light at an irradiance of 0.39 mW/mm^2^ for 4 hours. We validated MCI ROS induction by *nuo-1::Supernova* by observing increased fluorescence in the HyPer7 sensor (**Fig. S5**). Light-induced generation of MCI ROS by *nuo-1::Supernova* resulted in a modest but significant increase in sleep in wild-type animals (**Fig. 6B-C**). In addition, inducing MCI ROS prior to exposing animals to UVC irradiation, increased SIS more than UVC irradiation alone (**Fig. 6D**). These results suggest that mitochondrial ROS promotes sleep.

**Fig. 6.**
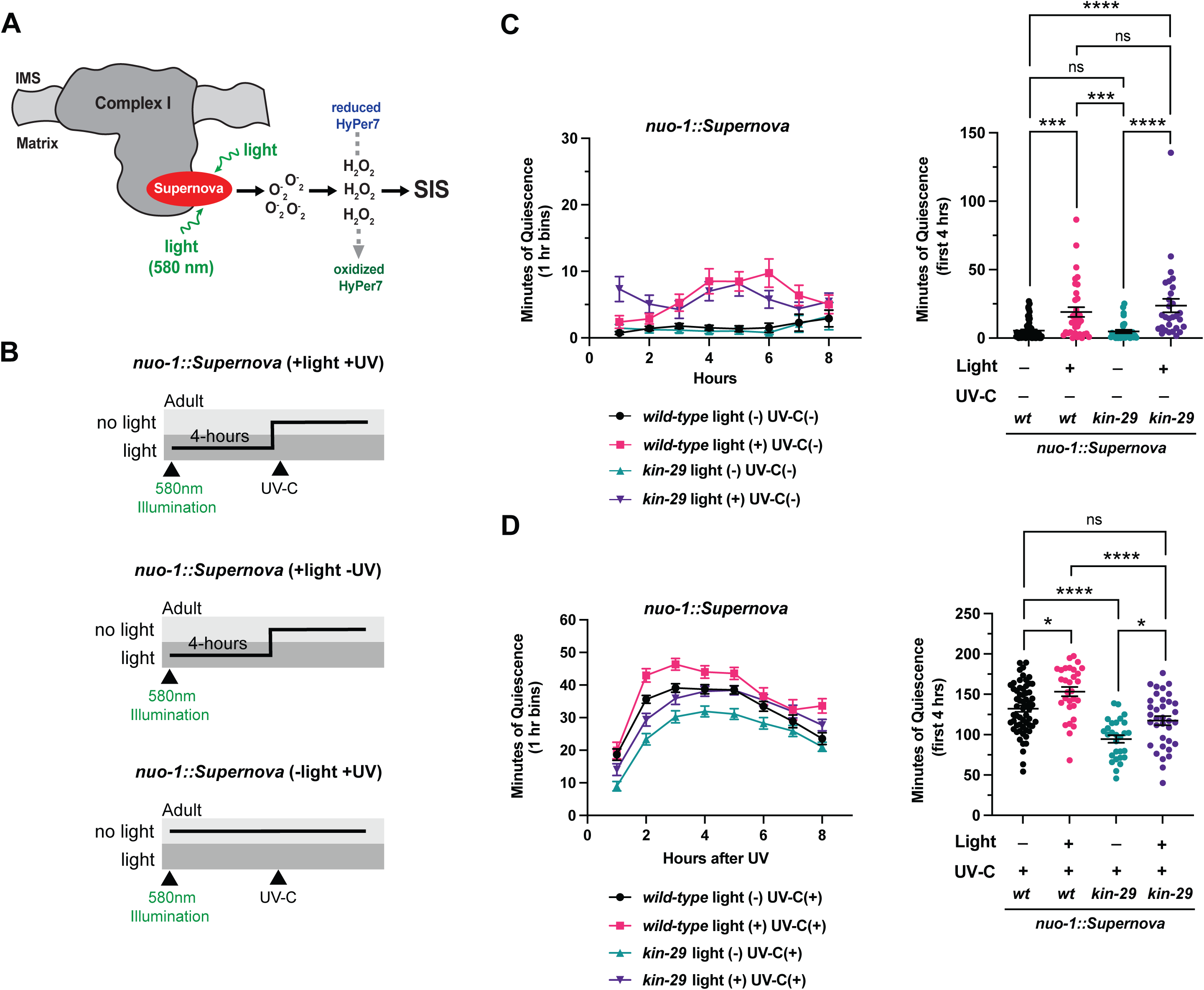
Mitochondrial ROS induction by Supernova promotes SIS and restores SIS in *kin-29* mutants. **(A)** Illustration of light-induced MCI ROS generation. Supernova is fused to the flavin mononucleotide (FMN) subunit, *nuo-1* (*31*). Upon light induction (580 nm), Supernova generates superoxide, which can then dismutate to H_2_O_2_ and is detected by the mitochondrial matrix-targeted HyPer7 biosensor. **(B)** Supernova activation protocols in age-matched young adults. *nuo-1::Supernova* (+light +UV) transgenic animals were exposed to light (580 nm, 0.39 mW/mm^2^) for 4 hours before being placed on a WorMotel chip and exposed to UVC (1,500 J/m^2^). *nuo-1::Supernova* (+light -UV) animals were exposed to light for 4 hours before being transferred to a WorMotel chip. *nuo-1::Supernova* (-light +UV) animals were exposed to UVC (1,500 J/m^2^) on a WorMotel chip. **(C)** Left: Time-course of body movement quiescence of wild-type and *kin-29(oy38)* mutants in response to Supernova activation by light (580 nm, 0.39 mW/mm^2^). Right panel: Minutes of movement quiescence during the first 4-hours post Supernova activation by light in wild-type (*wt*) and *kin-29(oy38)* mutants. Statistical significance was assessed using Kruskal-Wallis with Dunn multiple-comparisons test; \*\*\*\**P*<0.0001, \*\*\**P*<0.001; ns, not significant. **(D)** Left: Time-course of body movement quiescence of wild-type (*wt*) and *kin-29(oy38)* mutants in response to Supernova activation by light (580 nm, 0.39 mW/mm^2^) paired with UVC irradiation (1,500 J/m^2^). Right panel: Minutes of movement quiescence during the first 4-hours post Supernova activation by light (580 nm, 0.39 mW/mm^2^) in wild-type (*wt*) and *kin-29(oy38)* mutants. Statistical significance was assessed using one-way ANOVA with Tukey multiple-comparisons test; \*\*\*\**P*<0.0001, \**P*<0.05; ns, not significant. **(C-D)** n≥28 animals for each genotype after Supernova activation light and/or after UVC exposure. Data are represented as the mean ± SEM of 3 trials for each genotype and/or condition.

We next assessed whether light-induction of MCI ROS prior to somnogenic UVC exposure would rescue sleep in *kin-29* mutants, which exhibit low mitochondrial ROS levels. Light-induction of MCI ROS in *kin-29* mutants carrying the *nuo-1::Supernova* in the absence of UVC irradiation was sufficient to increase SIS to levels comparable to wild-type (**Fig. 6C**). Moreover, light-induction of MCI ROS in *kin-29* mutants with *nuo-1::Supernova* prior to UVC irradiation partially restored SIS in *kin*-*29* mutants to wild-type levels (**Fig. 6D).** When combined with UVC exposure, MCI ROS induction did not change the shape of the sleep curve; instead it increased the total amount of SIS in both wild-type and *kin-29* mutants. These results suggest that the role of mitochondrial ROS in promoting SIS is to modulate the amplitude of the sleep response.

KIN-29 acts in glutamatergic sensory neurons (defined by expression of the gene *odr-4*) to promote *C. elegans* sleep (*8*). These *odr-4*+ neurons act prior to activation of the central sleep promoting neurons ALA and RIS (*8*). However, the site of action of ROS to affect sleep remains unclear. To assess whether ROS modulation of sleep acts through the canonical SIS promoting ALA neurons, we introduced the *nuo-1::Supernova* transgene into the *ceh-17(np1)* loss-of-function mutant background; these mutants are impaired in the function of the sleep-promoting ALA neuron (*18, 21, 23, 43, 44*) (**Fig. 7A**). Light-induction of MCI ROS in *ceh-17(np1)* mutants with the *nuo-1::Supernova* transgene failed to induce sleep (**Fig. 7A**). Together, these results suggest that mitochondrial ROS signals within or upstream of the ALA sleep-promoting neuron.

**Fig. 7.**
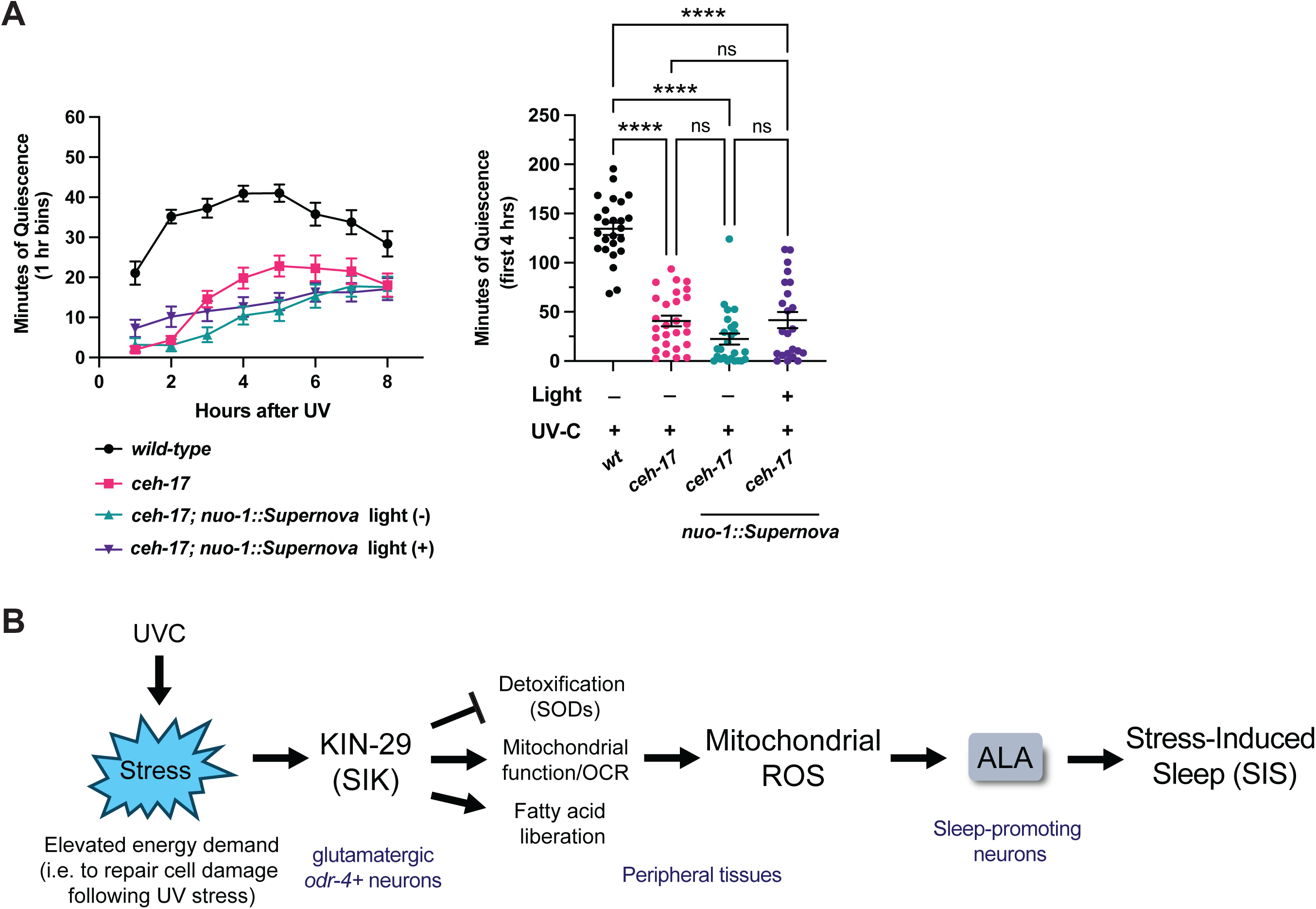
Mitochondrial ROS signals upstream of the ALA sleep-promoting neuron. **(A)** Left: Time-course of body movement quiescence of wild-type and *ceh-17(np1)* mutants with or without Supernova activation by light (580 nm, 0.39 mW/mm^2^). Right panel: Minutes of movement quiescence during the first 4-hours with or without Supernova activation by light in wild-type (*wt*) and *ceh-17(np1)* mutants. Statistical significance was assessed using a Kruskal-Wallis with Dunn multiple-comparisons test; \*\*\*\**P*<0.0001; ns, not significant. n≥24 animals for each genotype after UVC exposure with or without Supernova activation by light. Data are represented as the mean ± SEM of 3 trials for each genotype and/or condition. **(B)** Proposed model for KIN-29/SIK coupling of metabolic state to stress-induced sleep (SIS) through regulation of sleep-promoting mitochondrial ROS. During stress, such as UVC exposure, KIN-29/SIK promotes a transient increase in mitochondrial ROS and sleep. In the absence of KIN-29 function, there are three possible reasons for reduced ROS (and hence reduced sleep): (1) reduced fatty acid liberation via lipolysis, (2) reduced mitochondrial function, and (3) increased ROS toxification by SODs. Consistent with this model, *kin-29* mutants accumulate excess fat, β-oxidation is required for normal SIS, and increased fat mobilization partially restores SIS in *kin-29* mutants (*8*). KIN-29/SIK functions in 12 pairs of glutamatergic neurons defined by *odr-4* expression, upstream of ALA (and RIS) activation (*8*) to promote mitochondrial redox signaling and ROS production, which may in turn activate these sleep-promoting neurons to induce SIS. SIK, salt-inducible kinase, OCR, oxygen consumption rate.

## DISCUSSION

In this study, we identify mitochondrial reactive oxygen species (ROS) as sleep-promoting signals, and demonstrate that generation of this signal requires the salt- inducible kinase KIN-29, the *C. elegans* homolog of mammalian SIK3 (*8, 45*). Reduced ROS levels in *kin-29* mutants can be explained by reduced oxidative phosphorylation and upregulation of superoxide dismutase enzymes.

Once exposed to stress, *kin-29 mutants* fail to mount a mitochondrial ROS increase that coincides with peak SIS following UVC-induced stress. Importantly, restoring ROS optogenetically is sufficient to partially restore sleep in *kin-29* mutants, suggesting mitochondrial ROS acts as a downstream effector of KIN-29 in sleep regulation. Together, our findings extend prior work establishing KIN-29 as a key regulator of sleep-metabolism coupling (*8*) and place mitochondrial redox signaling downstream of this conserved SIK in the control of stress-induced sleep.

We show that mitochondrial ROS levels rise during SIS in wild-type animals, peaking during the period of maximal behavioral quiescence. This increase is delayed relative to UVC exposure, indicating that it is not simply a direct consequence of DNA damage or acute oxidative insult. Instead, mitochondrial ROS accumulation coincides with sleep itself, suggesting that ROS functions as a physiological signal associated with the initiation and/or maintenance of the sleep state. In contrast, *kin-29* mutants fail to generate this mitochondrial ROS response despite exposure to the same somnogenic UVC stress. This deficit is evident both at baseline and during SIS and is observed using whole-animal ROS measurements as well as *in vivo* imaging of intestinal mitochondrial hydrogen peroxide using HyPer7 (*31, 32*). These findings indicate that KIN-29 is necessary for enabling a stress-evoked mitochondrial ROS signal that normally accompanies sleep. Rather than being a downstream consequence of sleep, mitochondrial ROS is part of the signaling cascade that promotes SIS.

Our transcriptomic and proteomic analyses provide a partial explanation for the reduced mitochondrial ROS observed in *kin-29* mutants. Loss of KIN-29 results in upregulation of stress-response and detoxification pathways at both the transcript and protein levels, including strong induction of the mitochondrial superoxide dismutase SOD-3 (*35, 39*). This expression profile is consistent with enhanced ROS clearance and aligns with the increased resistance of *kin-29* mutants to paraquat-induced oxidative stress. These findings suggest that KIN-29 normally limits antioxidant gene expression, thereby allowing mitochondrial ROS to increase during stress. In the absence of KIN-29, elevated antioxidant capacity suppresses this mitochondrial ROS signal, thereby reducing sleep during SIS. In this model, reduced sleep in *kin-29* mutants reflects an inability to generate or maintain a sleep-promoting mitochondrial ROS signal. Consistent with this interpretation, genetic disruption of mitochondrial SODs enhances SIS, and the strongest effects are observed when all five SOD genes are removed. The UV-dose dependence of this phenotype indicates that increased sleep is specifically linked to impaired ROS clearance during oxidative stress, rather than to nonspecific locomotor dysfunction. Together, these data support a model in which mitochondrial ROS acts as a graded sleep-promoting signal whose amplitude is regulated by antioxidant capacity.

Our optogenetic experiments using *nuo-1::Supernova* that generates superoxide to H₂O₂-mediated ROS in the region of complex I (MCI), suggest that mitochondrial ROS acts downstream of KIN-29 to promote sleep. In *kin-29* mutants, ROS levels are low because of both reduced production (due to reduced mitochondrial function) and enhanced degradation (by up-regulating SODs). Targeted induction of mitochondrial ROS using Supernova fused to the mitochondrial respiratory chain complex I (MC I) (*31*) is sufficient to increase sleep in wild-type animals and partially restore SIS in *kin-29* mutants. This sleep induction occurs both in the absence of exogenous stress and following UVC exposure, demonstrating that mitochondrial ROS can bypass the requirement for KIN-29.

Supernova-mediated ROS induction increases the total amount of sleep without altering the temporal structure of the sleep response, suggesting that mitochondrial ROS modulates the amplitude of sleep drive rather than defining the timing of SIS. This is consistent with models from other systems in which ROS accumulates in proportion to sleep pressure and gates sleep initiation or depth (*46*). Our results therefore place mitochondrial ROS as a somnogenic signal of sleep downstream of KIN-29 function.

KIN-29 acts, at least in part, in *odr-4*-expressing sensory neurons, which are predominantly glutamatergic and act upstream of the sleep-promoting neurons ALA and RIS (*8*). Our finding that Supernova-induced mitochondrial ROS is blocked in *ceh-17* loss-of-function mutants, which have defective SIS-promoting ALA function (*18, 23*), suggests that mitochondrial ROS acts upstream of ALA. How mitochondrial ROS levels influence sleep-promoting neurons in *C. elegans*, and in which tissues this occurs, remains an open question. One possibility is that redox-sensitive signaling molecules or neuroendocrine pathways convey metabolic stress from KIN-29 acting in *odr-4*-expressing neurons to ALA/RIS.

Another possible source of mitochondrial ROS relevant to sleep may originate from mitochondrial β-oxidation (*47*). *kin-29* mutants have elevated fat stores (*8*), consistent with impaired lipid mobilization. β-oxidation is a major source of mitochondrial ROS, as increased electron flux through the respiratory chain enhances superoxide generation. In *C. elegans*, inhibition of β-oxidation reduces SIS, whereas increased fat mobilization partially restores sleep in *kin-29* mutants (*8*). Together these observations support a model in which KIN-29 activity in *odr-4* sensory neurons promotes signaling to fat storage tissues, triggering the release of energy stored as triglycerides and increases mitochondrial metabolic flux. This increase in metabolic flux may generate sleep-promoting mitochondrial ROS. In this model, the reduced mitochondrial ROS observed in *kin-29* mutants could result from reduced β-oxidation-derived ROS production and impaired mitochondrial function, potentially compounded by enhanced ROS detoxification. Thus, KIN-29 may couple metabolic state to sleep by regulating both the production and clearance of mitochondrial ROS. By promoting fat utilization while limiting antioxidant capacity, KIN-29 could enable a transient rise in mitochondrial ROS during stress, thereby translating metabolic imbalance into sleep drive.

Our results add to a growing body of evidence across species indicating that ROS can act as sleep-promoting signaling molecules (*46*). In flies and mice, sleep deprivation causes ROS to accumulate in defined sleep-regulatory neurons (*10, 12*), and experimental manipulation of redox balance within fly sleep circuits bidirectionally alters sleep (*11*). In addition to neuronal effects, sleep deprivation studies have identified lethal ROS accumulation in the gut (*15*), suggesting that peripheral oxidative stress may also contribute to sleep need. Consistent with this idea, prolonged total sleep deprivation in mice can escalate into a cytokine-storm-like inflammatory state (*17*), underscoring the close relationship between oxidative stress, immune signaling, and sleep loss. These findings align with long-standing literature showing that cytokines such as IL-1β and TNF-α can regulate sleep (*48*).

Together, these observations raise the possibility that mitochondrial ROS feeds into a broader, conserved stress program in which oxidative state helps shape inflammatory or cytokine-like signals that influence sleep drive. In *C. elegans*, SISS-1 is a damage-responsive LET-23/EGFR ligand required for SIS (*21*). SISS-1 release depends on ADM-4, the *C. elegans* ortholog of ADAM17/TACE (*21*), a metalloprotease that cleaves transmembrane substrates including EGF-family ligands and other cytokines (*49*). ADAM17/TACE activity can be stimulated by oxidative and osmotic stresses in mammalian cells (*50, 51*). Thus, one possibility is that mitochondrial ROS acts upstream of ADM-4/ADAM17-dependent shedding of SISS-1, thereby promoting LET-23/EGFR-activation in the ALA/RIS sleep circuit. In this model, mitochondrial ROS serves as a mechanistic link between metabolic or cellular stress and activation of the sleep circuit. More broadly, ROS-regulated ligand shedding may represent a conserved cytokine- or cytokine-like release mechanism through which peripheral damage signals are translated into sleep drive.

Finally, our results have implications regarding the mechanism of the sleepy phenotype in SIK3 gain-of-function mice, which show both elevated sleep and elevated sleep drive (*2, 52*). If mammalian SIK3 is functioning similarly to KIN-29, then it is reasonable to propose that ROS levels are high in SIK3 *Sleepy* gain-of-function mutants. In addition, mitochondrial SOD enzymes may be suppressed in SIK3 gain-of-function mutants. These hypotheses, which are readily testable, are relevant to mechanisms of sleep homeostasis throughout phylogeny.

### Limitations

There are 2 types of sleep in *C. elegans*—developmentally timed sleep (DTS) and stress-induced sleep (SIS)—and they are regulated differently in parts. Our experiments involved only SIS. KIN-29 also regulates DTS (*2, 8*) but we here did not pursue experiments with DTS.

In conclusion, our findings suggest that sleep is not merely protective against oxidative stress, but can be actively promoted by mitochondrial redox signals that report cellular damage and metabolic stress. In this framework, sleep represents an adaptive behavioral output of mitochondrial signaling pathway, with KIN-29/SIK3 coupling metabolic flux to sleep drive. Identifying the molecular sensors that detect mitochondrial ROS within the *C. elegans* sleep circuit, and determining how these signals are integrated with other stress-response pathways are important future directions

## MATERIAL AND METHODS

### *C. elegans* strains and growth conditions

Worms were cultivated at 20°C under standard conditions on Nematode Growth Medium (NGM) with *Escherichia coli* OP50 as the food source (*53*). The wild-type strain used was *C. elegans* variety Bristol, strain N2 (*54*); other strains used in this study are listed in **Table S1**. Genotypes were confirmed by PCR (for example, identifying deletions) or by sequencing PCR products (for example, identifying single nucleotide changes).

### Measurement of ROS

ROS levels in whole worm lysates were measured using the probe H_2_DCFDA (Sigma, cat# D6883) as described (*25*). Worms were age-matched via a double bleaching hypochlorite method (*55*) and transferred onto NGM plates covered completely with a lawn of *E. coli* OP50. Young adults were rinsed off a plate through a 35 μm nylon filter mesh (Sefar, Cat #7050-1220-000-10), which passes bacteria but traps young adult worms, and pelleted. Worms were lysed by bead beating for five minutes using a Bullet Blender Pro Storm (Next Advance). Lysates were flash frozen in liquid nitrogen and stored in -80°C until analysis. Protein concentration of worm lysates was determined using the MicroBCA Protein Assay Kit (ThermoFisher), and lysates were normalized to 25 μg protein per sample. Normalized lysates were incubated with H_2_DCFDA at a final concentration of 25 μM for 4 hours at 37°C in a 96-well plate. Fluorescence was measured using a Synergy HT plate reader (excitation 485 nm, emission 525 nm). Background fluorescence from no-worm controls was subtracted and values were normalized accordingly.

### Paraquat survival assay

Acute paraquat (PQ) resistance was assayed as described previously (*26*). Day 1 adult worms cultivated in non-crowded conditions and with an abundance of food (*E. coli*) were transferred using a platinum wire into 96-well plates filled with M9 containing 100 mM PQ. Each well contained 5-7 worms. Worms were scored every hour for being alive or dead by touching them with a platinum wire. Animals that did not move in response to this prod were considered dead. Scorers were blinded to the genotypes of each well until experiments were completed.

### Measurement of oxygen consumption rate (OCR)

OCR in whole worms was measured in an Agilent Seahorse XF Pro Analyzer using the Seahorse XF Pro M FluxPak (Agilent, 103777-100). Worms were age-synchronized using alkaline bleach method (*53*) and left rocking for 12-hours to arrest at L1 before plating on an NGM plate. Worms were aged to the L4 stage on the day of the assay. One day before the start of the assay, the XF Pro analyzer machine’s heater was shutdown and the machine was cooled to room temperature (22-23°C). On the day of the assay, the worms are washed off the agar surface of their cultivation plate using S-Basal and placed into 15 ml conical tubes. They were rinsed 3 times with 15 ml of S-Basal and then concentrated to 1 ml of S-Basal. 10 μl of worm solution was placed on a glass slide three times and worms were counted and averaged to determine the number of worms per μl in each condition. Each solution was then diluted so that wells in an XF Pro M Cell Culture Microplate had 160 μl of 0.3 worms/μl in S-Basal. Six wells were designated as blanks for the experiment only containing S-Basal. The worms were then imaged using a stereo microscope so that they could be later counted using ImageJ. OCR was measured using an Fe96 sensor cartridge with 4 cycles of basal readings, 7 cycles of 25 μM FCCP (Sigma Aldrich, SML2959-1 ml), and 4 cycles of 50 mM NaN_3_ in S-Basal. Each cycle was defined as mix 2 min, wait 30 seconds, measure 2 min. Once the machine was finished running, wells were screened to determine if drugs were properly injected. Well readings were then normalized by both worm count and body size. Once the data was normalized, the average of each well’s maximal cycles (7 readings) and the average of the basal cycles (4 readings) were recorded for each strain. The maximum-to-basal OCR ratio was calculated using the median of the averages to divide the maximal value by the basal value. This was repeated for each biological replicate.

### Size normalization of *C. elegans*

To accurately record OCR measurements, a worm’s size must be taken into account as smaller body sizes can have a reduced respiration despite normal mitochondrial function (*56*). To size normalize the worms, L4 worms were mounted on 5% agarose pads, immobilized with 50 mM levamisole, and imaged at 5x on a Leica DM5500 compound microscope equipped with a Hamamatsu Orca II camera. Images were then uploaded to ImageJ, and the mean in µm was used to normalize the oxygen consumption rate.

### Assessment of body movement quiescence following UVC stress

Movement quiescence was measured using a 48-well (6 x 8) WorMotel (*57*). First, synchronized day-old adult animals were transferred from their cultivation plate to WorMotel wells filled with NGM agar and spread with a thin layer of *E. coli* OP50 bacteria. They were then exposed to 1,500 J/m^2^ UVC (254 nm) in a Spectrolinker XL-1500. Immediately following UVC exposure, the WorMotel was placed in an uncovered plastic 10-cm Petri dish with moistened Kimwipes to maintain humidity. Lids were treated with a thin layer of 20% TWEEN® 20 to prevent condensation. The Petri dish was sealed to gas by stretching a sliver of Parafilm and imaged under dark-field illumination by red LED lighting. Images were captured every 10 seconds for the duration of the recording (8 hours) and analyzed with a custom MATLAB script (*58*). The quantification of movement was determined via pixel subtraction and an animal was considered quiescent if there was no pixel change between images (*59*). Worms that left the field of view and did not return as well as worms that burrowed into the agar were censored in the analysis (on average less than 20% for each assay). Trials were deemed useable if wild-type animals had 130 ± 25 minutes of quiescence during the first four hours post-UVC exposure.

### Total RNA collection and extraction

Animals were age-synchronized by alkaline hypochlorite treatment and collected eggs were hatched overnight at 20°C in 1x M9 buffer. L1 larvae were then plated onto NGM plates seeded with 10x concentrated *E. coli* OP50. Young adults were collected by quickly rinsing populations (less then 10 minutes) through a 35 μm nylon filter mesh (Sefar, Cat #7050-1220-000-10) with 1x M9 buffer, and flash frozen in liquid nitrogen. A 250 μl mixture of animals (worm pellet) in a 1x M9 buffer were then transferred into green RINO tubes (Next Advance) and resuspended with 750 μl of TRIzol LS reagent (ThermoFisher Scientific, Cat #10296028). Worms were immediately lysed by bead beating them for 5 min using a Bullet Blender Pro Storm (Next Advance). Total RNA was extracted using the PureLink RNA mini-kit (Ambion Cat#12183018A), followed by a DNAse I treatment according to the manufacturer’s protocol. Extracted RNA quality was quantified and assessed using an Agilent TapeStation 4150.

### RNA sequencing and analysis

One ug total RNA was extracted from five independent biological replicates from wild-type and *kin-29(oy38)* genotypes. Libraries were prepared and sequenced on an Illumina NovaSeq 6000 (Novogene) to generate 150bp paired-end reads. FASTQ files were trimmed using cutadapt(trim-galore) v4.4. Reads were mapped using STAR v2.7.11a to the reference UCSC genome ce11/WBcel235. Differential expression analysis was performed using DESeq2 v1.34. Raw FASTQ files from the RNA-seq data were deposited into the NCBI Sequence Read Archive (TBD). Individual accession numbers are listed in **Table S2**.

### Protein collection for mass spectrometry

Animals were age-synchronized by hypochlorite treatment and collected eggs were hatched overnight at 20°C in 1x M9 buffer. L1 larvae were then plated onto NGM plates seeded with 10x concentrated *E. coli* OP50. Young adults were collected by quickly rinsing populations through a 35μm nylon filter mesh (Sefar, Cat #7050-1220-000-10) with water. Animals were resuspended in RIPA lysis buffer (Pierce, 89901) containing protease inhibitors (Pierce, A32953), pelleted, lysed by bead beating, and flash frozen in liquid nitrogen.

### Protein digestion and mass spectrometry

Protein extraction and LC-MS/MS was performed by the IDeA National Resource for Quantitative Proteomics center for quantitative mass spectrometry. Proteins were extracted from five independent biological replicates from wild-type and *kin-29(oy38)* genotypes. Total protein from sonicated lysates was reduced, alkylated, and purified via chloroform/methanol extraction. Digestion was carried out with sequencing grade trypsin and LysC (Promega). Afterward proteins were labeled using the tandem mass tag 10-plex isobaric label reagent set (Thermo). Labeled peptides were separated into 46 fractions on a 100 x 1.0 mm Acquity BEH C18 column (Waters) using an UltiMate 3000 UHPLC system (ThermoScientific) with a 50 min gradient from 99:1 to 60:40 buffer A:B ratio under basic pH conditions (buffer A= 0.1% formic acid, 0.5% acetonitrile; buffer B = 0.1% formic acid, 99.9% acetonitrile), then consolidated into 18 super-fractions. Each super-fraction was then further separated by reverse phase XSelect CSH C18 2.5 um resin (Waters) on an in-line 150 x 0.075 mm column using an UltiMate 3000 RSLCnano system (ThermoScientific). A 75 min gradient was used to elute peptides from 98:2 to 60:40 buffer A:B ratio. Eluted peptides were ionized by electrospray (2.4 kV) followed by mass spectrometric analysis on an Orbitrap Eclipse Tribrid mass spectrometer (ThermoScientific) using multi-notch MS3 parameters. MS data were acquired using the FTMS analyzer in top-speed profile mode at a resolution of 120,000 over a range of 375 to 1500 m/z. Following CID activation with normalized collision energy of 31.0, MS/MS data were acquired using the ion trap analyzer in centroid mode and normal mass range. Using synchronous precursor selection, up to 10 MS/MS precursors were selected for HCD activation with normalized collision energy of 55.0, followed by acquisition of MS3 reporter ion data using the FTMS analyzer in profile mode at a resolution of 50,000 over a range of 100-500 m/z.

Data were processed by the Proteomics Center using MaxQuant software v2.2 against the *C. elegans* UniProt database (ID UP000001940) to identify proteins and quantify reporter ions. The following parameters were used: parent ion tolerance of 3 ppm, a fragment ion tolerance of 0.5 Da, a reporter ion tolerance of 0.001 Da, trypsin/P enzyme with 2 missed cleavages, variable modifications including oxidation on M, Acetyl on Protein N-terminus, and fixed modification of Carbamidomethyl on the C-terminus. Identified proteins with less than a 1% FDR were accepted. The Protein Prophet algorithm was used to assign protein probabilities. MS3 reporter ion intensities were normalized using ProteiNorm. Data was normalized using VSN and analyzed using Linear Models for Microarray data (limma) with empirical Bayes (eBayes) smoothing to the standard errors.

### Principle component analysis (PCA) and enrichment analysis

PCA was performed on feature counts of each RNA seq sample in R using the ade4 package (*60*). For proteomics, normalized mass spectrometry reads provided by the IDeA National Resource for Quantitative Proteomics center were used as input for PCA analysis using the same R software and packages. WormCat 2.0 was used to categorize enriched genes and proteins in both omics comparisons. The heat map was generated using the ComplexHeatmap package in R (*61*).

### Microscopy and fluorescence imaging

For the analysis of *Psod-3*::GFP expression, a Leica DM5500 compound microscope equipped with epifluorescence and a Hamamatsu CCD-camera was used to image mounted worms. Exposure time and camera settings were kept constant across samples. The mean pixel intensity of GFP fluorescence was quantified using Volocity software (version 6.3) in adult worms.

For HyPer7 imaging, worms carrying the intestinal mitochondrial-targeted HyPer7 (*Peft-1::2xCox8a::HyPer7*) (*31*) were age-matched by a double bleaching hypochlorite method (*55*) and selected for bright pharyngeal HyPer7 expression. Young adults were placed in 2 µL droplets of 0.10 μm immobilization beads (Cat# 00876-15) on 6% agarose pads in. For awake and SIS experiments, images of worm intestines were obtained as the ratio between excitation at 400 and 500 nm, with emission at 520 nm sequentially on a Leica THUNDER microscope. All images were acquired at the same exposure conditions. Signal was quantified as the ratio of fluorescence intensity changes with respect to the baseline using ImageJ (version 2.16.0) software.

### ROS induction by SuperNova

Worms were illuminated using a light-emitting diode (LED) system emitting 540-600 nm wavelength through a light guide (GYX module, 540-600nm, X-Cite LED1; Excelitas). To generate mitochondrial ROS by Supernova, worms carrying an integrated *nuo-1::Supernova* transgene (*31*) were exposed to light (580 nm, 0.39 mW/mm^2^) for 4 hours on 3.5 mm NGM plates freshly seeded with a thin layer of *E. coli* OP50 prior to transferring worms into the WorMotel. Immediately after light illumination, body movement quiescence with or without UVC irradiation was measured as described above. Irradiance was determined using an optical power meter (843R, Newport Corporation) and thermopile sensor (919P-020-12; Newport Corporation).

### Statistical analysis

Data was first subjected to a Shapiro-Wilk or D’Agostino-Pearson normality test (alpha = 0.05, *P* > 0.05). If a normality test was passed, then a parametric statistical test was performed. If a normality was not passed, then a non-parametric statistical test was used. Statistical comparisons made include the unpaired Welch’s *t*-test (parametric, 2 groups) or the unpaired Mann-Whitney test (non-parametric, 2 groups). For testing between three or more groups, we used one-way ANOVA (parametric) or Kruskal-Wallis (non-parametric) with post hoc multiple comparison correction. For other comparisons, statistical tests were performed as indicated in the figure legends. Statistical values of *P* ≤ 0.05 were considered significant. Data were visualized using Graphpad Prism (Version 10.6) software.

## Acknowledgments

We thank Andrew Wojtovich for the *nuo-1::Supernova* and HyPer7 transgenic strains, and Tyler G. O’Hara for assistance and training in the use of the Seahorse analyzer. Strains were provided by the *Caenorhabditis* Genetics Center (CGC), which is funded by NIH Office of Research Infrastructure Programs (P40 OD010440). We thank the IDeA National Resource for Quantitative Proteomics for performing mass spectrometry on our samples and providing guidance in experimental set-up. We thank the Cellular and Molecular Imaging Core facility of the COBRE Integrative Neuroscience Center at the University of Nevada for providing equipment and resources.

## Data, code and materials availability

Worm strains are available upon request from the authors. All data necessary for confirming the conclusions are present within the article, figures and tables. Raw FASTQ files from the RNA-sequencing data are deposited into the NCBI Sequence Read Archive (TBD) and NCBI Gene Expression Omnibus (GEO) (TBD). The mass spectrometry proteomics raw data are deposited to the ProteomeXchange consortium via the proteomics identification database partner repository (TBD).

