## Supplemental Figures for "KIN-29 SIK regulates stress-induced sleep through mitochondrial redox signaling"

### Supplementary Materials

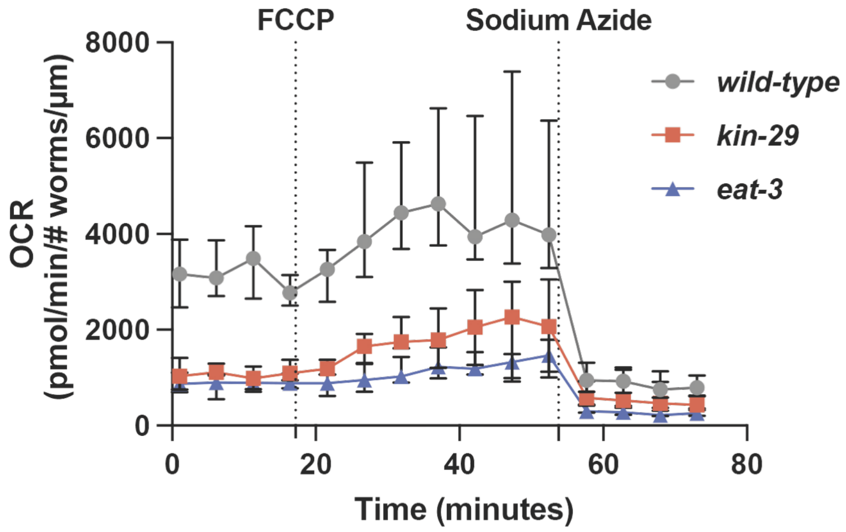


**Fig. S1. OCR measurements in *kin-29* mutants compared to wild-type and *eat-3* mutants**

Representative replicate showing the Oxygen Consumption Rate (OCR) measurements of wild-type (*wt*), *kin-29(oy38)*, and *eat-3(ad426)* mutant L4s treated with the uncoupler FCCP (25 uM) and the complex IV inhibitor $NaN_3$ (50 mM) in a timeline of 75 minutes using a Seahorse XF Analyzer. OCR readings were normalized by size and number of worms per well for each genotype. Data is represented by the median with the 95% confidence interval.

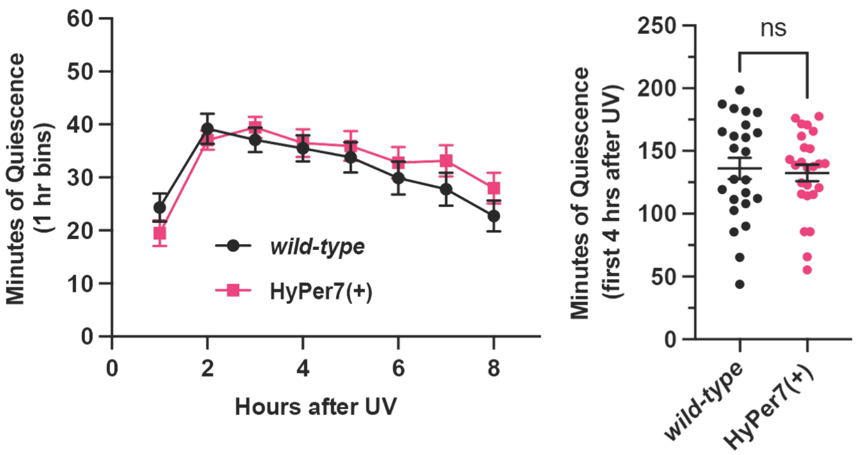


**Fig. S2. HyPer7 transgenic animals exhibit a wild-type SIS phenotype**

Time-course of body movement quiescence measured hourly over an 8-hour period (left graph) and minutes of quiescence during the first 4-hours (right graph) after UVC exposure in *Peft-3::2xCox8a::HyPer7* transgenic animals, HyPer7(+), compared with wild-type (*wt*) controls. n≥24 animals for each genotype after UVC exposure (1,500 J/m$^2$); 3 independent trials for each genotype. Data are represented as the mean ± SEM. Statistical significance was assessed using a two-tailed Mann-Whitney; ns, not significant.

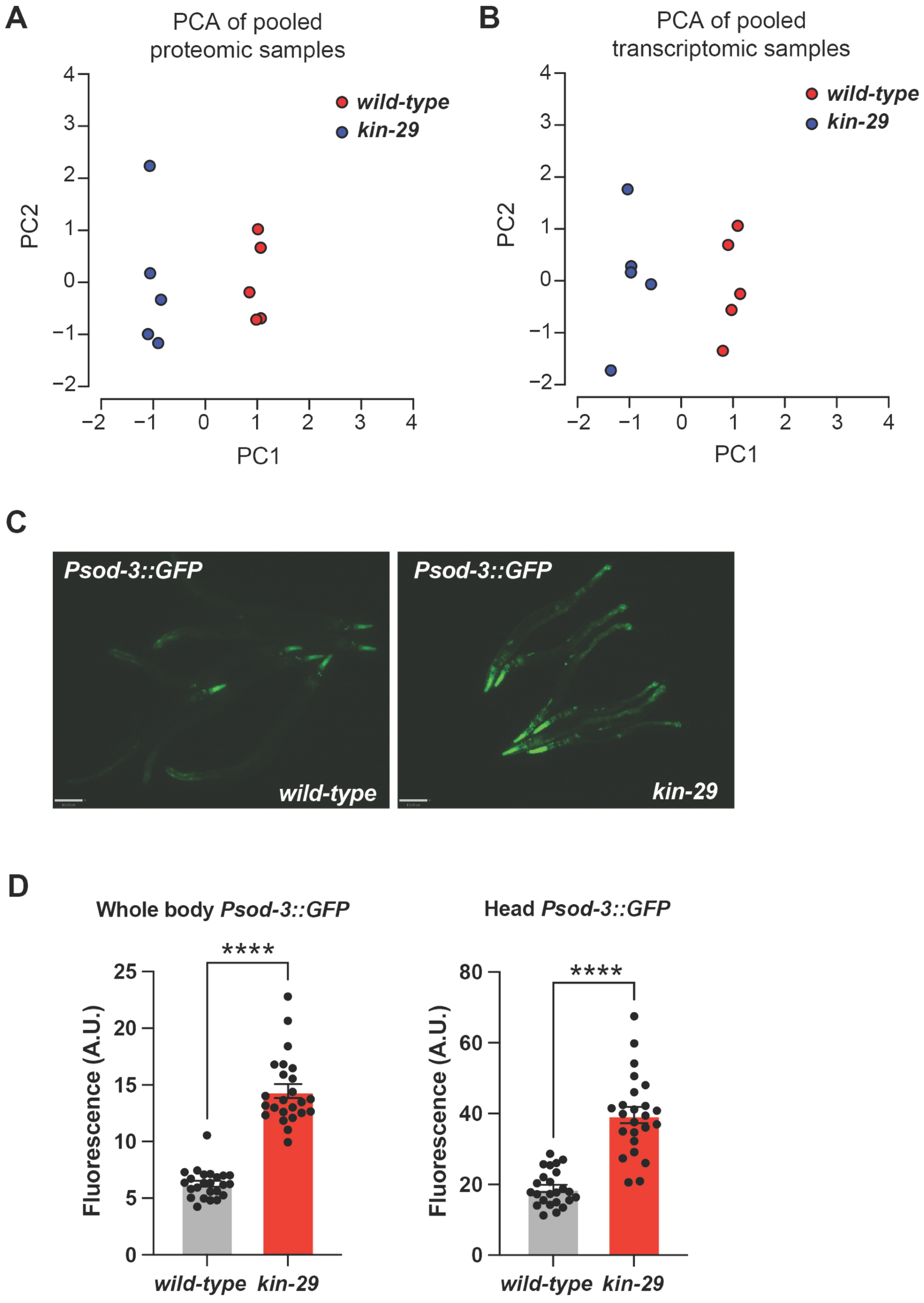
A
PCA of pooled proteomic samples
wild-type
kin-29
PC2
PC1
4
3
2
1
0
−1
−2
−2
−1
0
1
2
3
4
B
PCA of pooled transcriptomic samples
wild-type
kin-29
PC2
PC1
4
3
2
1
0
−1
−2
−2
−1
0
1
2
3
4
C
Psod-3::GFP
wild-type
Psod-3::GFP
kin-29
D
Whole body Psod-3::GFP
****
Fluorescence (A.U.)
25
20
15
10
5
0
wild-type
kin-29
Head Psod-3::GFP
****
Fluorescence (A.U.)
80
60
40
20
0
wild-type
kin-29

**Fig. S3. PCA analysis and *sod-3* expression between wild-type and *kin-29* mutants.**

**(A)** Principle Component Analysis (PCA) over two dimensions (PC1 and PC2) for RNA-Seq (left graph) and LC-MS/MS datasets in wild-type (blue color) vs *kin-29(oy38)* mutants (red color). The five biological replicates for each genotypes are well clustered. **(B)** *sod-3* expression increases in *kin-29(oy38)* mutants compared to wild-type. Shown are images of 5-6 whole animals expressing an integrated *Psod-3::GFP* transgene. Scale bar is 110 μm. **(C)** Quantification of *sod-3* expression in the head or whole body of animals. n≥20 animals for each genotype. ****$P$<0.0001 by a two-tailed Mann-Whitney test.

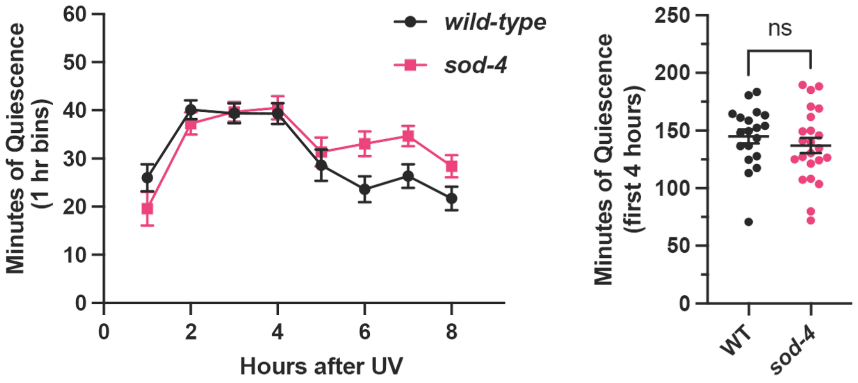


**Fig. S4.** ***sod-4*** **mutants exhibit a wild-type SIS phenotype.**

Time-course of body movement quiescence measured hourly over an 8-hour period (left graph) and minutes of quiescence during the first 4-hours (right graph) after UVC exposure for *sod-4(gk101)* mutants compared to wild-type (*wt*) controls. n≥19 animals for each genotype after UVC exposure (1,500 J/$m^2$); 3 independent trials for each genotype. Data are represented as the mean ± SEM. Statistical significance was assessed using a two-tailed Mann-Whitney; ns, not significant.

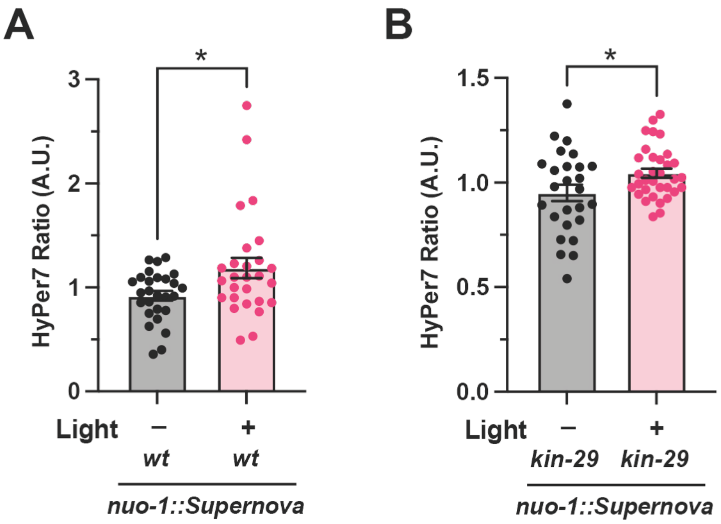


**Fig. S5. HyPer7 transgenic animals detect increased mitochondrial ROS following Supernova induction.**

Quantification of fluorescence of HyPer7 during wakefulness in wild-type (*wt*) and *kin-29(oy38)* mutant animals carrying both the *nuo-1::Supernova* and HyPer7 (*Peft-3::2xCox8a::HyPer7*) transgenes. **(A)** Relative HyPer7 intensity ratio (500/400 nm) in wild-type (*wt*) measured in response to Supernova activation by light (580 nm, 0.39 mW/mm$^2$). Data are represented as the mean ± SEM. n≥27 animals for each condition. *$P$<0.05 by a two-tailed Mann-Whitney test. **(B)** Relative HyPer7 intensity ratio (500/400 nm) in *kin-29(oy38)* mutants measured in response to Supernova activation by light (580 nm, 0.39 mW/mm$^2$). Data are represented as the mean ± SEM. n≥25 animals for each condition. *$P$<0.05 by a Welch's $t$-test.

**Table S1. (separate file)**

Strains used in this study

**Table S2. (separate file)**

RNA-Seq read statistics

**Table S3. (separate file)**

Differentially expressed genes in *kin-29* mutants

**Table S4. (separate file)**

Differentially expressed proteins in *kin-29* mutants

**Table S5. (separate file)**

Gene set enrichment analysis with WormCat 2.0
